# Stomach-brain synchrony binds neural representations of the body in a novel, delayed-connectivity resting-state network

**DOI:** 10.1101/217604

**Authors:** Ignacio Rebollo, Anne-Dominique Devauchelle, Benoît Béranger, Catherine Tallon-Baudry

## Abstract

Resting-state networks offer a unique window into the brain’s functional architecture, but their characterization remains limited to instantaneous connectivity thus far. Here, we describe a novel resting-state network based on the delayed connectivity between the brain and the slow electrical rhythm (0.05 Hz) generated in the stomach. The gastric network cuts across classical resting-state networks with little overlap with autonomic regulation areas. This network is composed of regions with convergent functional properties involved in mapping bodily space through touch, action or vision, as well as mapping external space in bodily coordinates. The network is characterized by a precise temporal sequence of activations within a gastric cycle, beginning with somato-motor cortices and ending with the extrastriate body area and dorsal precuneus. Our results demonstrate that canonical resting-state networks based on instantaneous connectivity represent only one of the possible partitions of the brain into coherent networks based on temporal dynamics.

## Introduction

The parsing of the brain into resting-state networks (RSNs) has been widely exploited to study the brain’s functional architecture in health and disease (1). With long time scales, RSNs closely match the anatomical backbone of the brain (2–4). With short time scales (~10-100 s), spontaneous brain activity is characterized by the emergence and dissolution of network patterns encompassing and extending classical RSN topologies (5, 6) with rich temporal trajectories (7). Temporal trajectories indicate the existence of delays between regions, whereas the methods most often used to parse brain activity into functional networks (seed-based correlation and independent component analysis) make the implicit assumption that RSNs are characterized by instantaneous or zero delay connectivity. Therefore, we analyzed delayed connectivity in resting-state BOLD signals using techniques widely used in electrophysiological studies of large-scale brain dynamics (8) that quantify the stability of temporal delays between time series.

More specifically, we studied the delayed coupling between resting-state brain activity and a visceral organ, the stomach. The stomach continuously produces a slow electrical rhythm (0.05 Hz, one cycle every 20 seconds) that can be non-invasively measured (electrogastrogram (EGG) (9)). The gastric basal rhythm is continuously (10) and intrinsically (11) generated in the stomach wall by a network of specialized cells, the interstitial cells of Cajal (12), which form synapse-like connections not only with gastric smooth muscle but also with afferent sensory neurons (13). The stomach is an interesting candidate for large-scale brain coordination for several reasons. First, visceral inputs can reach a number of cortical targets (14, 15). Second, gastric frequency (~0.05 Hz) falls within the range of BOLD fluctuations that are used to define RSNs and are free from known cardiac and respiratory artifacts (16). Finally, the amplitude of alpha rhythm, the dominant rhythm in the human brain at rest, depends on the phase of gastric rhythm (17).

We simultaneously recorded brain activity with fMRI and stomach activity with EGG (Fig. 1a) in 30 human participants at rest with open eyes. We then determined the regions in which spontaneous fluctuations in brain activity were phase synchronized with gastric basal rhythm; we refer to these regions as the gastric network.

**Fig. 1.**
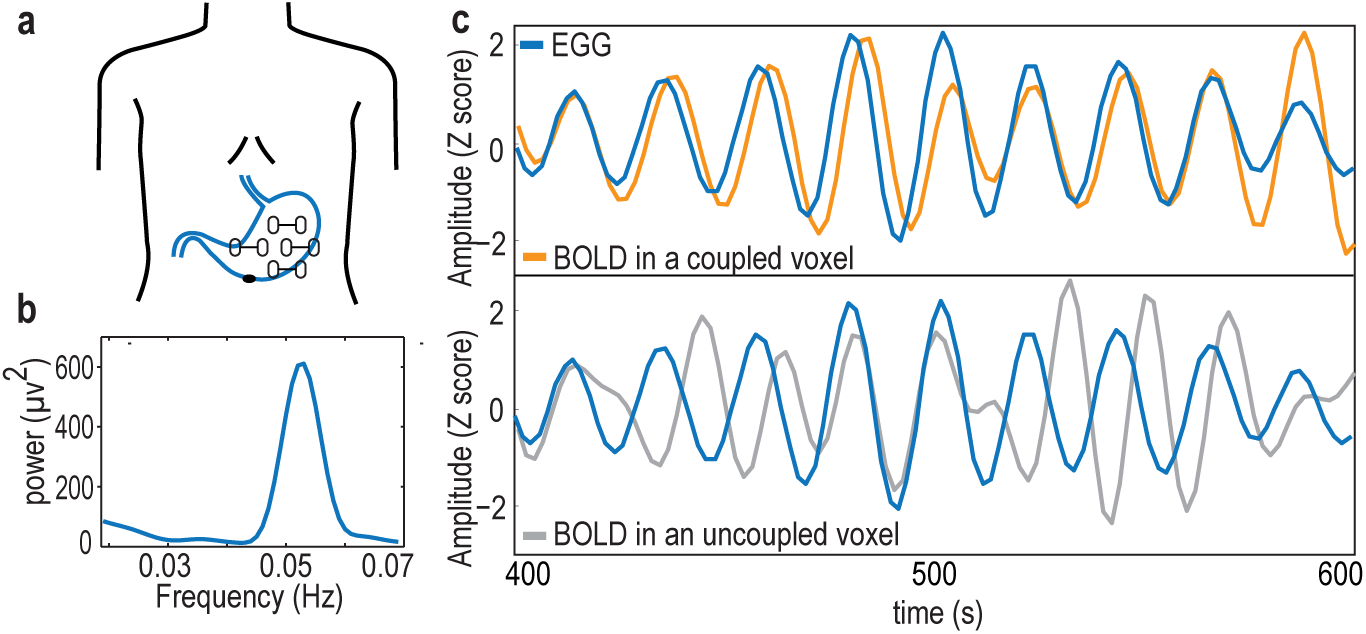
The electrogastrogram (EGG) and gastric-BOLD coupling. **a.** Bipolar electrode montage used to record the EGG. **b.** Example of an EGG spectrum in a single participant, with the typical spectral signature in the normogastric frequency range (0.033-0.066 Hz). **c.** Example of a 200-s time series of the EGG and BOLD signal filtered at the gastric peak frequency in an EGG-coupled voxel (top), characterized by a stable, non-zero phase relationship between EGG and BOLD, resulting in a high phase-locking value, and in an EGG-uncoupled voxel (bottom) in which no stable phase relationship between EEG and BOLD can be observed.

## Results

### EGG-BOLD phase coupling defines the gastric network

We first determined gastric frequency (Fig. 1b) in each participant as the frequency of the largest spectral peak within the normogastric range (0.033-0.066 Hz). The mean EGG peak frequency across the 30 participants was 0.047 Hz (± SD 0.003, range 0.041-0.053). EGG peak frequency measured inside and outside the scanner did not differ (EGG outside the scanner measured in 29 of the 30 participants, mean 0.046 Hz ± SD 0.006; two-sided paired t-test, t(28)=0.35, p=0.725 Bayes Factor<0.001, indicating decisive evidence for the null hypothesis).

In each participant and at each voxel, we quantified the degree of phase synchrony between the EGG signal and BOLD time series filtered around gastric frequency (Fig. 1c). We computed the phase-locking value (PLV) (8), a measure widely used in electrophysiology that varies between 0 when two time series show no consistent phase relationship (Fig. 1c, bottom panel) and 1 when two time series have a consistent phase relationship over time (Fig. 1c, upper panel). Importantly, the PLV is high for any lag between the time series as long as this lag is constant over time. In each participant and at each voxel, we estimated that the PLV that could be expected by chance from EGG signals that were shifted with respect to the BOLD time series. The empirical PLVs were then compared with chance-level PLVs using a cluster-based statistical procedure that intrinsically corrects for multiple comparisons (18). Significant phase coupling between the EGG and resting-state BOLD time series occurred in twelve nodes (voxel threshold p<0.01, two-sided paired t-test between the observed and chance PLV; cluster threshold corrected for multiple comparisons, Monte-Carlo p<0.05, two-sided).

The gastric network (Table 1, Fig. 2a) comprises the right primary somatosensory SIr, bilateral secondary somatosensory SII, medial wall motor regions (MWM, comprising the caudate cingulate motor zone (CCZ), posterior rostral cingulate motor zone (RCZp), and right supplementary motor area (SMA)), a region of the right occipito-temporal cortex overlapping the extrastriate body area (EBA), as well as nodes in the posterior cingulate sulcus, dorsal precuneus, occipital cortex (ventral and dorsal portions), retrosplenial cortex (RSC), and superior parieto-occipital sulcus. The average shared variance between the EGG and BOLD signals across participants, as estimated from squared coherence, ranged from 12% in the left anterior dorsal precuneus to 16.9% in the posterior cingulate sulcus (pCS) (Table 1). An analysis of covariance across nodes did not reveal significant links between gastric-BOLD coupling strength and gender (F(1, 28)=1.02, p=0.46), body mass index (BMI) (F(1, 28)=1.3, p=0.3) or trait anxiety scores (F(1, 28)=1.02, p=0.47).

**Table 1:**
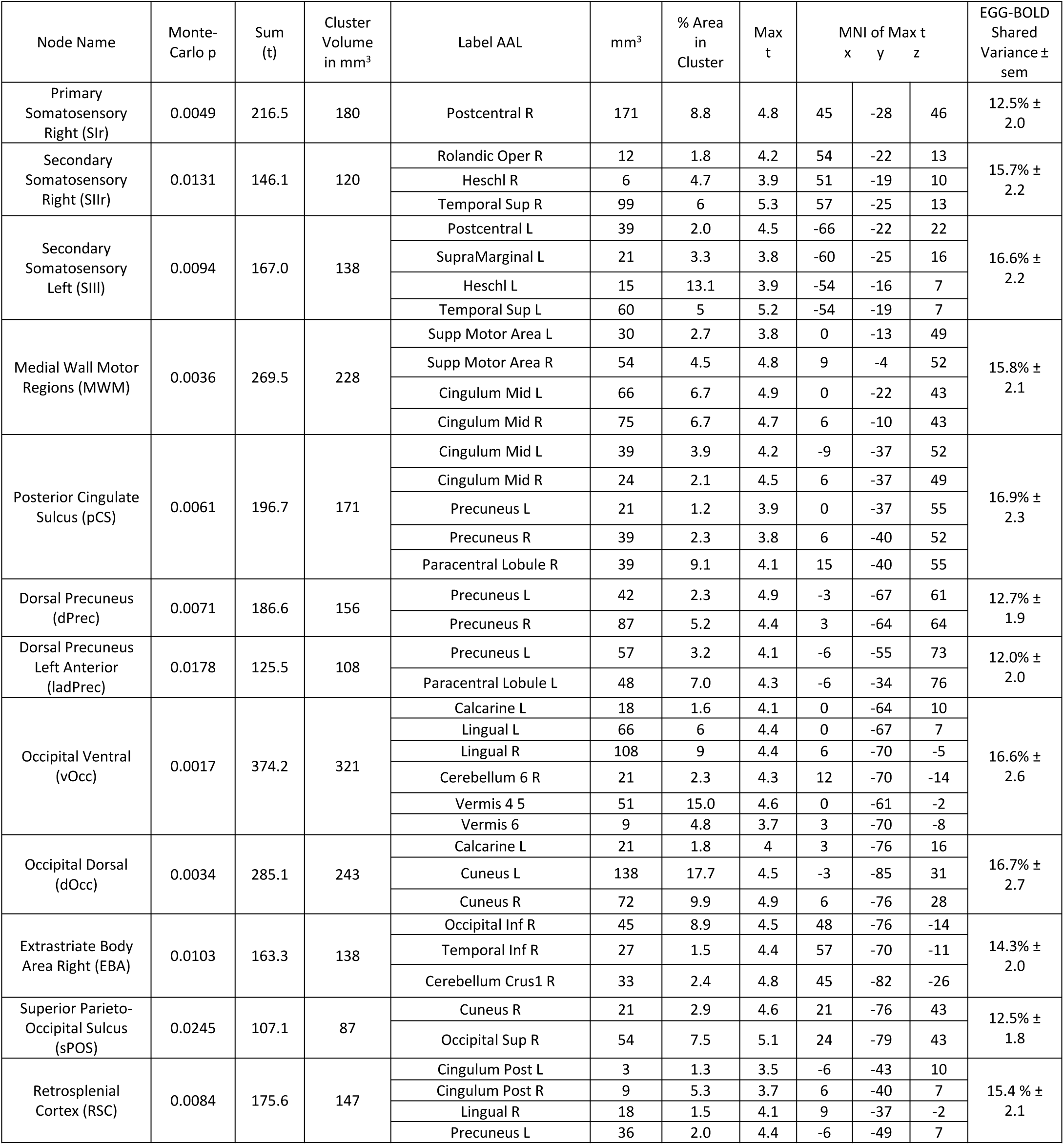
Description of the twelve nodes showing larger-than-chance gastric-BOLD phase coupling. AAL: Automated Anatomical labeling (58). MNI: Montreal Neurological Institute.

**Fig. 2.**
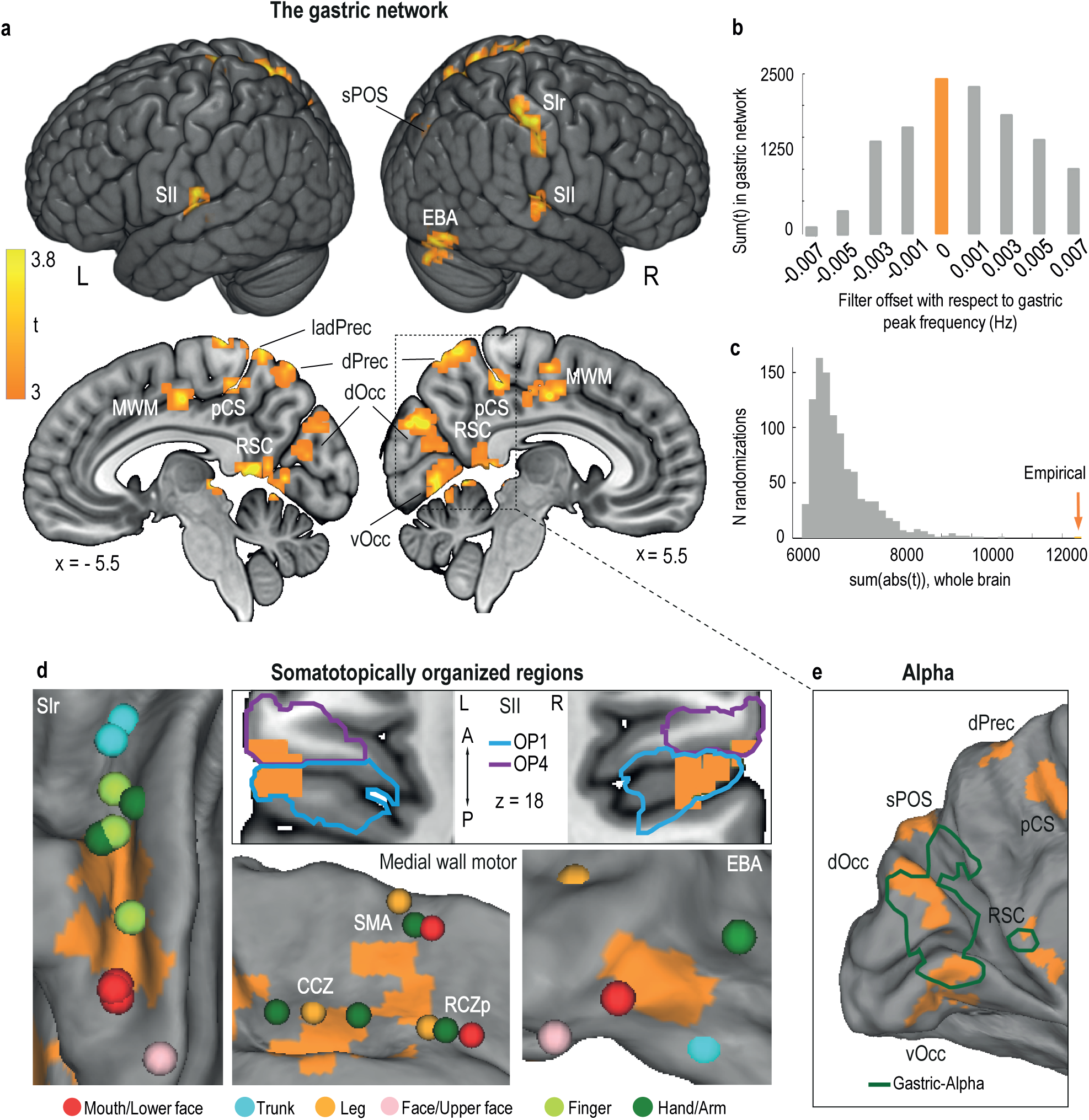
The gastric network. **a.** Regions significantly phase synchronized to gastric rhythm (N=30, voxel level threshold, p<0.01 two-sided; cluster level threshold, p<0.05, two-sided, corrected for multiple comparisons). **b.** Gastric-BOLD coupling is specific to gastric frequency. Summary statistics in the gastric network are maximal at the EGG peak frequency (orange) and decrease when offsetting the filter to higher or lower frequencies. **c.** Summary statistics distribution under the null hypothesis from 1000 surrogate datasets in which the EGG signal was time-shifted with respect to the BOLD signal. The empirical finding (orange arrow) falls well outside the null distribution. **d.** The gastric network (orange) comprises the following somatotopically organized regions: *primary somatosensory cortex* (*Panel SI,* with peak activations during stimulation of the trunk and hand (59), finger (60), face (61), and mouth, i.e., teeth (62), lips and tongue (63); *secondary somatosensory cortex* (*Panel SII*, cytoarchitectonic subdivisions of SII according to (22); OP1, parietal operculum 1 and OP4, parietal operculum 4, presented on a horizontal slice at z=18); *medial wall motor areas* (*Panel MWM*, with peak activations during movement (23) in the caudate cingulate zone (CCZ), posterior rostral cingulate zone (RCZp) and supplementary motor area (SMA)); and *extrastriate body area* (*Panel EBA* with peak activations during body part viewing (25), note that because of the visualization on an inflated cortex, the extension of the EBA node to the cerebellum is not visible). **e.** Regions in which the alpha and gastric rhythms are coupled (green, modified from (17)). Abbreviations are the same as those in Table 1.

### Controls: gastric frequency specificity, false-positive rate, and head micromovements

To assess the robustness of the gastric network, we ran several controls. First, we verified that EGG-BOLD coupling was specific to gastric frequency. We filtered both EGG and BOLD time series at frequencies that were slightly offset from the peak gastric frequency of each participant and recomputed cluster statistics. Summary statistics (sum of the absolute t-values resulting from the paired t-test between empirical and chance-level PLV at each voxel, either summed across the whole brain or within the gastric network) decreased when shifting below or above the gastric peak frequency (Fig. 2b). This result indicates that the gastric network corresponds to BOLD fluctuations specifically occurring at gastric frequency.

Second, we estimated the likelihood of false positives with our statistical procedure. We randomly sampled surrogate datasets in which a random time shift was applied to the EGG of each participant a thousand times. Next, we tested whether any of those 1000 combinations would generate summary statistics as large as the original data when compared with the chance-level estimate we used to determine significantly coupled regions at the group level (Fig. 2c). This result was never observed, indicating that the probability of our results being a false positive is less than 0.001.

Finally, we investigated whether submillimeter head movements might have influenced the results. We defined voxel motion susceptibility as the regression coefficient of head movement (19) from the BOLD time series. Coupling strength and voxel motion were unrelated (Fisher z-transformed Pearson correlation coefficients tested against zero, t(29)=-0.34, p=0.73; Bayes factor<0.001, indicating decisive evidence for the absence of a link between coupling strength and head movement). Stomach contractions might also lead to small head movements that could be phase locked to gastric rhythm. Although gastric rhythm is continuously produced even during fasting, it is larger during stomach contractions. Thus, we tested whether the effects we found were due to differences in EGG power (or frequency) across participants. We found no link between coupling strength (difference between empirical and chance PLV) in the 12 nodes and EGG power (ANCOVA, F(1, 28)=0.9, p=0.51; all Bayes factor<0.33, indicating substantial evidence for the null hypothesis) nor between coupling strength and EGG peak frequency (ANCOVA, F(1, 28)=1.6, p=0.17; Bayes Factor<0.33, indicating substantial evidence for the null hypothesis in nine of twelve nodes; Bayes Factor<1.3 in the three remaining nodes, indicating anecdotal or no evidence).

The gastric network is thus specific to individual gastric peak frequency, is highly unlikely to be a chance finding, and is not linked to spurious effects of head movement on the BOLD signal.

### The gastric network includes body maps associated with touch, action and vision

We then examined the areas comprising the gastric network in more detail. By definition, the gastric network is composed of regions with activity that co-fluctuates with gastric basal rhythm. Five nodes of the gastric network also share a common functional feature, somatotopic organization, as detailed in Fig. 2d.

The gastric network includes the following regions with a well-known body representation based on touch: the right primary somatosensory cortex in the hand and mouth region and bilateral secondary somatosensory cortices. We further quantified the overlap between the nodes of the gastric network and known cytoarchitectonic subdivisions of the somatosensory cortices (20, 21). The gastric network mostly overlapped with area 1 (60.2% of the SIr node) and to a lesser extent, with area 2 (13.1%) and area 3b (9.9%). The SII nodes of the gastric network overlapped with the secondary somatosensory cortices and more precisely with the somatotopically organized subdivisions of the parietal operculum OP1 and OP4 (22). The right SII node mostly overlapped with area OP1 (35.2% of the node), while the left SII node overlapped with both OP1 (21.7%) and OP4 (14.9%). Additionally, both left and right SII nodes extended more ventrally to the temporal cortex.

The gastric network also includes three medial wall motor regions (CCZ, RCZp, and SMA) that reveal their somatotopic organization when participants are required to move specific body parts (23). Note that gastric-BOLD coupling also included a more posterior area in the cingulate sulcus (pCS). Finally, the gastric network overlapped with the EBA, a region of the lateral occipital cortex activated when participants view images of body parts (24) with a clear somatotopic organization (25). The overlap between the gastric network and EBA occurred in the lower face region, which includes the mouth.

Thus, the gastric network overlaps with body maps classically associated with different modalities, including touch in somatosensory cortices, action in MWM and vision in the EBA.

### The gastric network includes regions involved in the generation of the alpha rhythm

Finally, we found gastric-BOLD coupling in the posterior bank of the parieto-occipital sulcus (dOcc and vOcc) and retrosplenial cortex. In a previous study using magneto-encephalography (17), the amplitude of the alpha rhythm in these regions was modulated by gastric phase (Fig. 2e).

### Marginal gastric-brain coupling in the insula and autonomic control networks

The insula is one region that receives visceral inputs (14, 15), but it did not appear to be significantly phase synchronized to the EGG using our whole-brain conservative statistical procedure. Thus, we performed post hoc region-of-interest analysis of the three insular subdivisions (anterior dorsal, anterior ventral, posterior (26)) in both hemispheres. Only the right posterior insula showed some indication of gastric-BOLD coupling across participants (empirical vs. chance-level PLV, paired t-test, two sided, t(29)=2.78, p=0.043, Bonferroni corrected; all other regions, p>0.21).

Brain-body interactions studies usually focus on brain regions that directly impact the bodily state via sympathetic or parasympathetic outputs. The overlap between the gastric network and autonomic regulation areas (27) was very limited (34 voxels, 4.7% of the gastric network) and confined to SIr and the anterior parts of SMA and RCZp (Fig. 3).

**Fig. 3.**
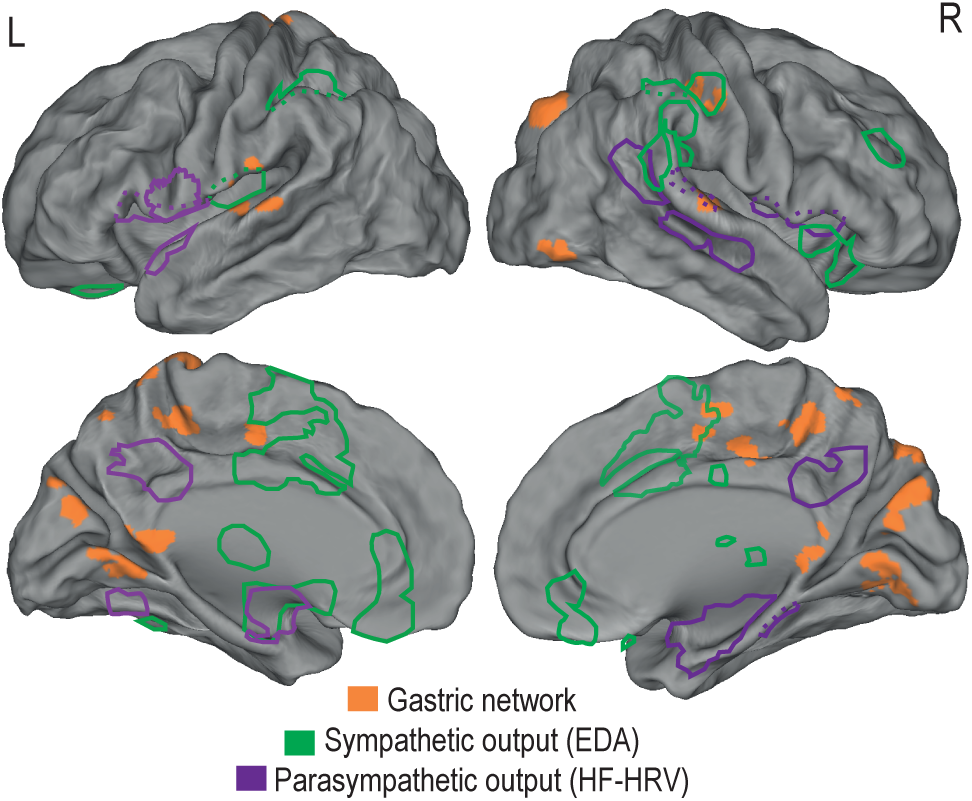
The gastric network is distinct from autonomic control regions. Meta-analytic activations (27) during tasks eliciting sympathetic (electrodermal activity, EDA, in green) and parasympathetic (high frequency heart rate variability, HF-HRV, in purple) responses are superimposed on the gastric network (orange). Dotted lines illustrate the extension of autonomic activity in the depth of the opercular and temporal regions.

### Temporal sequence within a gastric cycle and delayed connectivity between the nodes of the gastric network

In the different nodes of the gastric network, gastric-brain coupling occurred with different phase delays with respect to the gastric cycle. We analyzed between-participant phase-delay consistency and found temporal delays of ~3.3 seconds between the earliest nodes (somatosensory cortices) and latest nodes (dorsal precuneus and EBA) of the gastric network (Fig. 4a,b). The Watson-Williams test for circular data confirmed that different nodes of the gastric network were coupled to the gastric rhythm with different phase delays (F(11, 29)=5.22, p<10^−6^), indicating a precise temporal sequence of activations within each gastric cycle.

**Fig. 4.**
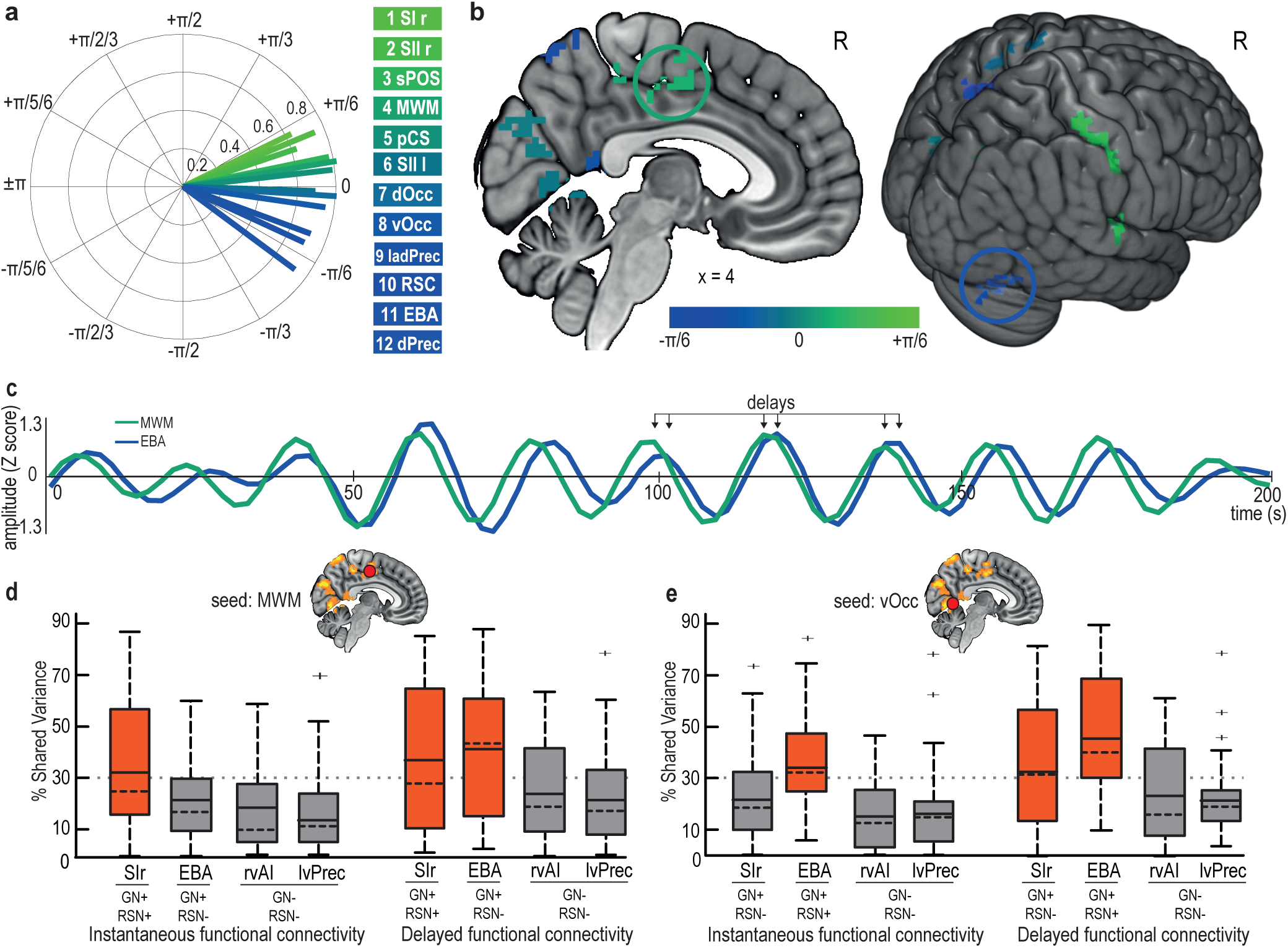
Early and late nodes of the gastric network are functionally connected but with a delay. **a.** Phase-delay consistency of each node of the gastric network. Vector length quantifies the phase consistency across participants, and vector angle indicates phase advance (green) or phase lag (blue) within the gastric network. Temporal delays can reach up to ~3.3 s (±π/6) between the earliest nodes (primary and secondary somatosensory cortices) and the latest nodes (dorsal precuneus and EBA) **b.** Group-averaged phase delays for each cluster in the gastric network. The two circled regions (EBA, blue; MWM, green) are illustrated in C. **c.** A 200-s BOLD time series in MWM and EBA in a single participant, showing phase consistency with delays. **d**. Group-level functional connectivity across all participants using MWM as a seed, either instantaneous (left) or delayed (right), with regions belonging or not to the gastric network (GN+/-, defined by delayed connectivity) and with regions belonging or not to the same resting-state network (RSN+/-, defined by instantaneous connectivity). Boxes are colored red when the mean FC exceeds 30%. The boxplot presents the mean (full line), median (dashed line), first and third quartiles (box), and extrema (whiskers) excluding outliers (+, defined as exceeding 1.5 interquartile ranges above the third quartile). **e.** Group-level functional connectivity across all participants using the vOcc as a seed. Abbreviations: EBA, extrastriate body area; MWM, medial wall motor regions; rSI, right primary somatosensory cortex; rVAI, right ventral anterior insula; lvPrec, left ventral precuneus.

Thus, each node of the gastric cycle appears to be characterized by a specific temporal delay with respect to gastric phase. These temporal delays were accompanied by delayed functional connectivity (FC) between the nodes of the gastric network.

We first illustrated this point with an example in a single participant (Fig. 4c), with two 200-s time series of the gastric network (MWM and EBA). The two time series systematically co-varied with a temporal delay. The existence of temporal delays between the nodes of the gastric network is one of the reasons why the gastric network could not be observed in prior studies. Indeed, fMRI RSN studies are typically based on measures of instantaneous FC, such as shared variance estimated from the squared Pearson correlation coefficient, which does not detect the temporally delayed interactions revealed here. These measures differ from delayed FC measures based on the consistency of phase delays over time, such as shared variance estimated from squared coherence. In the example illustrated in Fig. 4c, instantaneous FC between the two time series is 56%, whereas delayed FC is 86%. If we advance the timing of the medial wall time series by 2 s, instantaneous FC increases to 86%. This finding shows that the difference between the two FC estimates is due to temporal delays only.

We then estimated both instantaneous and delayed FC between all nodes of the gastric network in all participants. Delayed FC between gastric nodes (mean 40.8% ± SD 8%, ranging from 26.5% between the right primary somatosensory cortex and RSC, up to 63.9% between the ventral and dorsal occipital cortices) was systematically larger (paired t-test, t(29)=9.02, p<10^−10^) than instantaneous FC (mean 30.2% ± SD 11%, ranging from 9.3% between the dorsal precuneus and right SII, up to 61.2% between right and left SII). Next, we verified (Fig. 4d, e) that two regions belonging to both the gastric network and the same RSN (i.e., two regions of the gastric network with little temporal delay, such as MWM and SIr) would display large values of both delayed and instantaneous FC, whereas two regions belonging to the gastric network but not to the same classical RSN (i.e., two regions of the gastric network with a large temporal delay, such as MWM and EBA) would show large delayed FC and small instantaneous FC. Thus, in contrast to classical RSNs, the gastric network appears to be characterized by between-node delayed connectivity.

### Slow temporal fluctuations in gastric-BOLD coupling are associated with changes in BOLD amplitude and occur simultaneously in all nodes

Thus far, we have identified a sequence of activation that occurs at each gastric cycle, which characterizes gastric-BOLD coupling. We then investigated whether slow temporal fluctuations in the strength of gastric-BOLD coupling were accompanied by fluctuations in BOLD amplitude. As illustrated in Fig. 5a, we found that episodes of elevated gastric-BOLD synchronization corresponded to episodes of increased BOLD amplitude. Indeed, time-varying PLV and BOLD time series, computed in sliding time windows of 60 s (approximately 3 gastric cycles), were significantly correlated (Fisher z-transformed Pearson correlation coefficients t-tested against zero, Bonferroni corrected p<0.006 in all gastric nodes, mean r across nodes 0.18 STD=±0.02, ranging from 0.15 in MEM to 0.22 in SIIl).

**Fig. 5.**
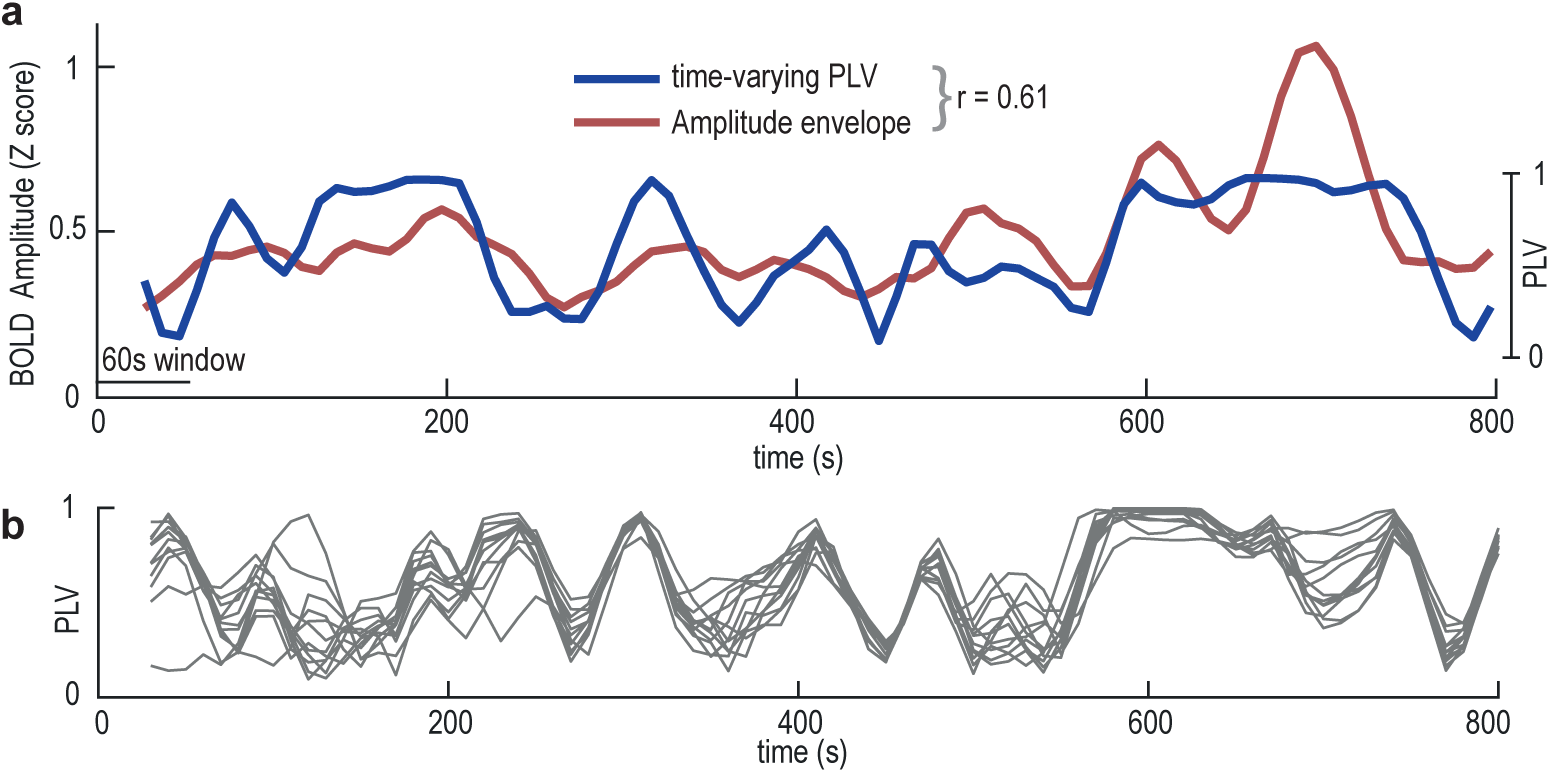
Dynamic fluctuations in EGG-BOLD coupling. **a**. Data from the superior parieto-occipital sulcus in a single participant showing the correlation (r=0.61) between the time-varying gastric-BOLD coupling (PLV, blue) and the amplitude envelope of the BOLD time series (red). **b.** Time-varying PLV at each node of the gastric network in the same participant showing a strong correlation across nodes (average r across all possible node pairs 0.35).

Next, we tested whether slow temporal fluctuations in gastric-BOLD synchronization occurred simultaneously or independently in the different nodes of the gastric network (Fig. 5b). We computed the correlation between time-varying PLVs for all possible node pairs in each participant and found that at the group level, this correlation was significantly positive (Fisher z-transformed average Pearson correlation coefficients against zero, t(29)=9.22, p<10^− 10^, mean r=0.129 ± STD 0.075, range across participants 0.02–0.35). To determine whether the overall pattern of synchronous fluctuations in gastric-BOLD coupling strength was driven by specific node pairs, we investigated correlations between node pairs. All node pairs but RSC-SIr, sPOS-RSC, sPOS-SIIr and sPOS-ladPrec showed a significant positive correlation at the group level (Fisher z-transformed Pearson correlation coefficients in all pairs tested against zero, Bonferroni corrected). The three nodes showing the largest covariation in time-varying PLV with the other nodes were dPrec, pCS and vOcc, and the three nodes showing the least covariation in time-varying PLV with the other nodes were RSC, SIr and sPOS.

## Discussion

Here, we reveal the existence of the gastric network, comprising brain regions with BOLD time series that are phase synchronized with gastric basal rhythm. Within the gastric network, approximately 15% of the BOLD variance is explained by gastric-BOLD phase synchrony. The gastric network cuts across classical RSNs and shows little overlap with autonomic control regions. A number of brain regions composing the gastric network have convergent functional properties involved in mapping bodily space through touch, action and vision. The network is characterized by a precise temporal sequence of activations within a gastric cycle, beginning with somato-motor cortices and ending with extra-striate body area and dorsal precuneus. This temporal sequence is accompanied by delayed functional connectivity between nodes of the gastric network, which explains why this RSN could not be identified with standard correlation methods that only capture instantaneous connectivity. Thus, our results suggest that canonical RSNs based on instantaneous connectivity represent only one of the possible partitions of the brain into coherent networks based on temporal dynamics.

### Neural origin of gastric-BOLD coupling

SIr, SII and medial wall motor regions likely receive direct gastric inputs. The stimulation of the splanchnic (spinal) nerve that innervates the stomach evokes responses in contralateral SI and bilateral SII in several mammals (28), and the spinothalamic tract was recently shown to target MWM in monkeys (29). Vagal stimulation can also evoke responses in somato-motor cortices (30). In addition, single neurons with convergent visceral and hand inputs have been observed in SI (31), in line with the overlap we observed between the gastric network and hand representation. SIr, SII and MWM are not only targeted by documented ascending gastric pathways, they are also early nodes of the gastric network, with a phase advance compared with that of other nodes. Thus, these areas could be the entry point of gastric afferences, relaying gastric information via cortico-cortical connectivity to other nodes such as the EBA, a late node of the gastric network.

Since regions receiving direct visceral inputs are also early nodes of the gastric network, the BOLD fluctuations locked to the gastric rhythm likely have a neural origin. An additional argument is that we found gastric-BOLD coupling in parieto-occipital regions with neural activity in the alpha range modulated by gastric phase (17). However, below, we examine the possibility that other non-neural mechanisms might contribute to gastric-BOLD coupling.

Artifactual BOLD fluctuations caused by head movements driven by stomach contractions seem unlikely. Indeed, gastric-BOLD coupling was neither related to head movement nor to EGG power that increases during stomach contractions. Another possibility is a vascular artifact. During digestion, gastric blood flow does indeed vary (32), but cerebral blood flow is unaltered (33). Artificial distension of the stomach can cause increases in peripheral blood pressure (34), but this peripheral increase is mostly due to the insertion of a bag catheter, not to its inflation (35). Finally, spontaneous fluctuations in blood pressure in humans occur at approximately 0.1 Hz (so-called Mayer waves), which is much faster than gastric rhythm. Thus, a vascular effect seems unlikely, and the hypothesis that activity in the gastric network is driven by neural activity in areas directly receiving ascending inputs that later propagates to other nodes of the gastric network appears more plausible.

### Gastric network and bodily space

The gastric network appears to be organized along a common functional principle related to bodily space. Gastric basal rhythm creates a functional link between body maps corresponding to different sensory modalities. In addition to the primary and secondary somatosensory cortices, the gastric network includes MWM (CCZ, RCZp and SMA) that are involved in motor preparation and display a clear somatotopic organization (23, 36). The gastric network also comprises the EBA, a functional region in the occipito-temporal cortex that selectively responds to visual images of the human body (24, 37) and is causally involved in body visual recognition (38), with a fine topographical organization (25). The EBA is not purely visual since it is also activated when participants move or imagine body parts without visual feedback (39). Body maps associated with touch, action or vision thus appear to be functionally coupled via the stomach, an organ that cannot be easily touched, moved or seen.

In addition to body maps associated with touch, action and vision, the gastric network comprises regions that play a role in mapping the external space in bodily coordinates, namely, the right superior parieto-occipital sulcus, dorsal precuneus and RSC. The superior parieto-occipital sulcus region is a visuo-motor area that encodes visual stimuli in bodily coordinates during action (40). The dorsal precuneus and RSC both implement the integration of information into an egocentric reference frame, a key basic mechanism involved in many different cognitive processes (41, 42) that is, by definition, centered on the body.

Thus, in the brain at rest, gastric rhythm appears to bind together distributed maps encoded in bodily coordinates into a coherent functional system.

### The gastric network is a novel resting-state network

RSNs have been defined as segregated functional systems that show synchronous fluctuations during rest (1). The gastric network, albeit distinct from classical RSNs as well as from the autonomic modulation areas (27), falls under this definition. In terms of dynamics, the gastric network is defined by its phase synchronization with the stomach and its delayed connectivity between nodes. Functionally, the gastric network anchors different body-centered representations within a single functional system. The gastric network can thus be considered a novel RSN that could not be previously observed due to methodological reasons.

As opposed to classical RSNs, the gastric network is characterized by delayed connectivity, with temporal delays that can extend to several seconds but are stable over time and captured by coherence and phase synchrony. Delays are an intrinsic characteristic of brain dynamics unfolding in anatomically connected networks (43) and pervasive even at the timescale of BOLD signal fluctuations (7). Canonical RSNs based on instantaneous connectivity represent only one of the possible partitions of the brain into coherent networks based on temporal dynamics. Therefore, we propose the addition of delayed connectivity to the operational definition of RSNs.

## Methods

### Experimental procedure

#### Participants

Thirty-four right-handed human participants took part in this study. All volunteers were interviewed by a physician to ensure the following inclusion criteria: absence of digestive, psychiatric or neurological disorders; BMI between 18 and 25; and compatibility with MRI recordings. Participants received a monetary reward and provided written informed consent for participation in the experiment. The study was approved by the ethics committee *Comité de Protection des Personnes Ile de France III*. All participants fasted for at least 90 minutes before the recordings. Data from four participants were excluded. Two were excluded because coughing artifacts caused excessive head movement during acquisition and corrupted the EGG data, and two were excluded because their EGG spectrum did not show a clear peak that could allow us to identify the frequency of their gastric rhythm. A total of 30 participants (mean age 24.2 ± SD 3.31, 15 females, mean BMI 21.48 ± SD 1.91) were included in the analysis described below.

#### MRI data acquisition

MRI was performed at 3 Tesla using a Siemens MAGNETOM Verio scanner (Siemens, Germany) with a 32-channel phased-array head coil. The resting-state scan lasted 900 s during which participants were instructed to lay still and fixate on a bull’s eye on a gray background. A functional MRI time series of 450 volumes was acquired with an echo-planar imaging (EPI) sequence and the following acquisition parameters: TR=2000 ms, TE=24 ms, flip angle=78°, FOV=204 mm, and acquisition matrix=68x68x40 (voxel size=3x3x3 mm^3^). Each volume comprised 40 contiguous axial slices covering the entire brain. High resolution T1-weighted structural MRI scans of the brain were acquired for anatomic reference after the functional sequence using a 3D gradient-echo sequence (TE=1.99 ms, TR=5000 ms, TI-1=700 ms/TI-2=2500 ms, flip angle-1=4°/flip angle-2=5°, bandwidth=240 Hz/pixel, acquisition matrix=240×256×224, and isometric voxel size=1.0 mm^3^). The anatomical sequence duration was 11 minutes 17 s. Cushions were used to minimize head motion during the scan.

#### Physiological signal acquisition

Physiological signals were simultaneously recorded during functional MRI acquisition using MRI compatible equipment. The electrogastrogram (EGG) and electrocardiogram (ECG) were acquired using bipolar electrodes connected to a BrainAmp amplifier (Brain products, Germany) placed between the legs of participants; the electrodes received a trigger signaling the beginning of each MRI volume. EGG was acquired at a sampling rate of 5000 Hz and a resolution of 0.5 μV/bit with a low-pass filter of 1000 Hz and no high-pass filter (DC recordings). ECG was acquired at a sampling rate of 5000 Hz and a resolution of 10 μV/bit with a low-pass filter of 1000 Hz and a high-pass filter of 0.016 Hz. Eye position and pupil diameter were recorded with an EYELINK 1000 (SR Research, Canada) and simultaneously sent to BrainAmp amplifiers.

The skin of participants was rubbed and cleaned with alcohol to remove dead skin, and electrolyte gel was applied improve the signal-to-noise ratio. The EGG was recorded via four bipolar electrodes placed in three rows over the abdomen, with the negative derivation placed 4 cm to the left of the positive one. Fig. 1a shows the electrode placement scheme. The midpoint between the xyphoid process and umbilicus was identified, and the first electrode pair was set 2 cm below this area, with the negative derivation set at the point below the rib cage closest to the left mid-clavicular line. Another electrode pair was set 2 cm above the umbilicus and aligned with the first electrode pair. The positive derivation of the third pair was set in the center of the square formed by electrode pairs one and two. The positive derivation of the fourth electrode pair was centered on the line traversing the xyphoid process and umbilicus at the same level as the third electrode. The ground electrode was placed below the lower left costal margin. The ECG was acquired using three bipolar electrodes that shared the same negative derivation, set at the third intercostal space. The positive derivations were set at the fifth intercostal space and separated by 4 cm.

Electrophysiological data were collected during fMRI data acquisition, as well as at least 30 s before and after. In addition, to rule out the possibility that the scanner pulse and B0 magnetic field could distort the frequency content of the EGG, a second EGG acquisition with an 8-min duration was performed after the acquisition of the MRI scans, with the participant positioned outside the tunnel of the scanner. Paired sample t-test was then performed to compare the peak frequencies obtained for each participant inside the scanner with those obtained outside the scanner for the same channels. This control analysis was run on 29 participants due to corrupted data in the EGG recordings outside the scanner tunnel in one participant.

#### MRI preprocessing

Brain imaging data were preprocessed using Matlab (Matlab 2013b, MathWorks, Inc., United States) and the Statistical Parametric Mapping toolbox (SPM 8, Wellcome Department of Imaging Neuroscience, University College London, U.K.). Images of each individual participant were corrected for slice timing and motion with 6 movement parameters (3 rotations and 3 translations). Two participants who moved more than 3 mm during the functional scan were excluded from the study. Each participant’s structural image was normalized to Montreal Neurological Institute (MNI) space of 152 participants’ average T1 template provided by SPM with affine registration followed by nonlinear transformation (44, 45). The normalization parameters determined for the structural volume were then applied to the corresponding functional images. The functional volumes were spatially smoothed with a 3 mm^3^ full-width half-maximum (FWHM) Gaussian kernel. The time series of voxels inside the brain, as determined using a SPM a priori mask, were subjected to the following preprocessing steps using the FieldTrip toolbox (46) (Donders Institute for Brain, Cognition and Behaviour, Radboud University Nijmegen, the Netherlands. See <u>http://www.ru.nl/neuroimaging/fieldtrip</u>, release 01/09/2014). Linear and quadratic trends from each voxel’s timeseries were removed by fitting and regressing basis functions, and we bandpass filtered the BOLD time series between 0.01 and 0.1 Hz using a 4^th^ order Butterworth infinite impulse response filter. A correction for cerebrospinal fluid motion was obtained by regressing out the timeseries of a 9-mm diameter sphere located in the fourth ventricle (MNI coordinates of the center of the sphere [0 -46 -32]).

#### EGG preprocessing

Data analysis was performed using the FieldTrip toolbox. Data were low-pass filtered below 5 Hz to avoid aliasing and downsampled from 5000 Hz to 10 Hz. To identify the EGG peak frequency (0.033-0.066 Hz) for each participant, we computed the spectral density estimate at each EGG channel over the 900 s of an EGG signal acquired during the fMRI scan using Welch’s method on 200-s time windows with 150-s overlap. Spectral peak identification was based on the following criteria: peaking power and sharpness of the peak. Two participants were excluded from further analysis at this stage because their spectral peak was unclear, with a power smaller than 15 μV^2^. The two criteria coincided in twenty participants. In ten participants, we used the second most powerful channel because the spectral peak was sharper. Data from the selected EGG channel was then bandpass filtered to isolate the signal related to gastric basal rhythm (linear phase finite impulse response filter, designed with Matlab function FIR2, centered at ECG peaking frequency, filter width ± 0.015 Hz, filter order of 5). Data were filtered in the forward and backward directions to avoid phase distortions and downsampled to the sampling rate of the BOLD acquisition (0.5 Hz). Filtered data included 30 s before and after the beginning and end of MRI data acquisition to minimize ringing effects.

MR gradient artifacts affect the electrophysiological signal down to approximately 10 Hz, which is far above EGG frequency (~0.05 Hz). Thus, no specific artifact gradient procedure was necessary. We further checked that EEG frequency inside and outside the scanner did not differ (see Results).

### Data analysis

#### Quantification of Gastric-BOLD phase synchrony

The BOLD signals of all brain voxels were bandpass filtered with the same filter parameters as the ones used for the EGG preprocessing. The first and last 15 volumes (30 s) were discarded from both the BOLD and EGG time series. The updated duration of the fMRI and EGG signals in which the rest of the analysis was performed was 840 s. The Hilbert transform was applied to the BOLD and EGG time series to derive the instantaneous phases of the signals. The PLV(8) was computed as the absolute value of the time average difference in the angle between the phases of the EGG and each voxel across time (Equation 1).

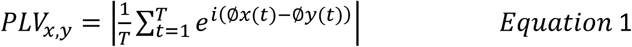

where T is the number of time samples, and x and y are the two time series. The PLV measures phase synchrony irrespective of temporal delays and amplitude fluctuations and is bounded between 0 (no synchrony) and 1 (perfect synchrony). The PLV was first assessed over the whole duration of the recording. In a second step, we computed the time-varying PLV in a 60-s time window shifted by 10 s.

Statistical procedure for determining regions showing significant gastric-BOLD coupling at the group level

We employed a two-step statistical procedure adapted from a previous work (17). We first estimated chance-level gastric-BOLD coupling at each voxel and in each participant. We then used group-level statistics to determine regions in which gastric-BOLD coupling was greater than chance.

We first estimated the chance-level PLV at each voxel for each participant. We created surrogate datasets in which the phase relationship between the EGG and BOLD time series was disrupted by offsetting the EGG time series with respect to the BOLD time series. In practice, the EGG time series was shifted by a time interval of at least ±60 s (i.e., approximately 3 cycles of the gastric rhythm) with respect to the BOLD time series. Data at the end of the recording were wrapped to the beginning. Given the 420 samples in the BOLD time series, this procedure generated 360 surrogate datasets from which we could compute the distribution of the chance-level PLV for each voxel in each participant. The chance-level PLV was defined as the median value of the chance-level PLV distribution for each voxel and participant. We defined coupling strength as the difference between the empirical PLV and chance-level PLV.

In a second step, we tested whether the empirical PLV differed from the chance-level PLV across participants. We used a cluster-based permutation procedure (18), as implemented in FieldTrip (46), that extracts clusters of voxels showing significant differences at the group level while intrinsically correcting for multiple comparisons. This non-parametric method is exempt from the high rate of false positives associated with the Gaussian shape assumption often present in fMRI studies (47). The procedure consists of comparisons between the empirical PLV and chance-level PLV across participants using t-tests at each voxel. Candidate clusters are formed by neighboring voxels exceeding the first-level t-threshold (p<0.01, two-sided). Each candidate cluster is characterized by the sum of the t-values in the voxels defining the cluster. To determine the sum of t-values that could obtained by chance, we computed a cluster statistics distribution under the null hypothesis by randomly shuffling the labels “empirical” and “chance level” 10.000 times and applied the clustering procedure. At each permutation, we retained the largest positive and smallest negative summary statistics obtained by chance across all voxels and thus built the distribution of cluster statistics under the null hypothesis and assessed the empirical clusters for significance. Because the maximal values across the whole brain are retained to build the distribution under the null hypothesis, this method intrinsically corrects for multiple comparisons. Clusters are characterized by their summary statistics (sum(abs(t))) and Monte-Carlo p value. Clusters with a Monte-Carlo p value<0.05 (two-sided, corrected for multiple comparisons) were considered significant and are reported in the Results section as nodes of the gastric network.

### Quantification of gastric-bold shared variance

To estimate the amount of variance in the BOLD signal that could be accounted for by gastric coupling, we computed the squared coherence coefficient between the EGG and average BOLD time course across all voxels in each significant cluster using FieldTrip software. The coherence coefficient measures phase and amplitude consistency across time and is a frequency domain analog of the cross-correlation coefficient in the temporal domain. Therefore, its squared value can be interpreted as the amount of shared variance between two signals at a certain frequency (48). First, we estimated the frequency spectrum of the full-band (0.01-0.1 Hz) EGG and BOLD signals (Welch method on a 120-s time window with 20-s overlap). We then computed the coherence coefficient between the spectrum of each participant’s EGG and each cluster’s time series at gastric frequency (*ω*)42T as the absolute value of the product of the amplitudes (A) of the signals and their phase (φ) difference averaged across time windows (t) and normalized by the square root of the product of their squared amplitudes averaged across time windows.

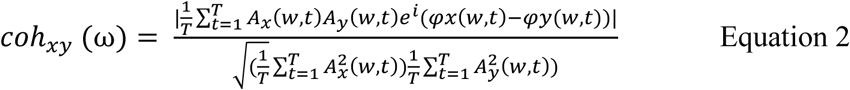

The coherence coefficient was then squared and averaged across participants such that the final group value represented the shared variance between the EGG and each cluster BOLD activity at the normogastric peak.

### Between-participant phase-delay consistency

To quantify temporal delays in the gastric network, we ran group-level analysis on the gastric-BOLD phase-locking angle. In each participant, we first computed a mean BOLD time series per node by averaging the voxel time series in each significant cluster. We then computed the relative phase-locking angle ϕ_k_ relative of the node *k* between the node time series *x* and the EGG *y* using equation 3, where ϕ_k_ relative corresponds to the phase-locking angle ϕ_k_ of node k with respect to the EGG minus the average angle across all nodes. ϕ_k_ relative thus quantifies the phase advance or lag of each node relative to the gastric network.

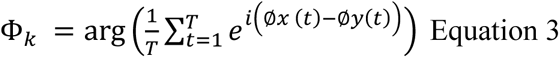

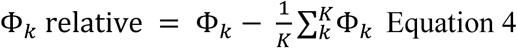

Between-participant phase-delay consistency was then obtained at each node by averaging the unit vectors of the relative phase-locking angles across *P* participants using equation number 5.

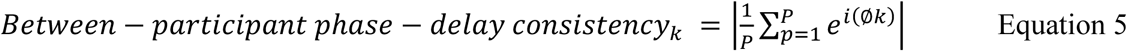

To determine whether there were significant differences across the angle of gastric network clusters, we submitted the values of each node and participant’s relative phase-locking angle to Watson-Williams test, a circular analog of one-way ANOVA for circular data, using the circstat Matlab toolbox (Berens et al., 2009).

### Functional connectivity: correlation and coherence

FC was defined as shared variance and computed using either the squared Pearson correlation coefficient or squared coherence. We computed the Pearson correlation between the bandpass-filtered BOLD time series (gastric peaking frequency ± 0.015 Hz) averaged across voxels in each gastric node, as well as in two control regions outside the gastric network, the right ventral precuneus and right ventral insula. The ventral precuneus, a core node of the default network, was defined using a 9 mm^3^ ROI centered in the coordinates provided by (50) (MNI x=-5 y=-52.5 z=41). The right ventral insula ROI was provided by the parcellation performed by (26).

To compute coherence between BOLD time series, we first estimated the frequency spectrum of the full-band (0.01-0.1 Hz) BOLD time series using the Welch method with 36 time windows of 120 s with 20-s overlap. We then computed coherence using the FieldTrip implementation of equation number 2 and used the squared coherence at the gastric peak frequency of each participant as an estimate of shared variance.

### Bayes factor

Bayesian statistics on correlation coefficients were computed and interpreted according to (51) and (52). Regarding the specific test of an absence of effect of voxel motion susceptibility on coupling strength (H0), submillimeter voxel motion was estimated as in (19), and H1 was modeled as the minimum effect size required to detect a significant difference from zero, given one-sample t-test of 29 degrees of freedom on a normal distribution with a mean of 0 and a standard deviation of 1. The same method was used to test for the absence of a difference between the EGG peak frequency measured inside and outside the scanner.

### Anatomical and functional overlays and meta-analysis

Functional group-level images were overlaid on a 3D rendering of the MNI template using MRIcroGL (<u>https://www.nitrc.org/frs/?group_id=889</u>, June 2015). The results from the literature were converted when necessary from Talairach coordinates to MNI coordinates using the nonlinear transform provided by (http://imaging.mrc-cbu.cam.ac.uk/imaging/MniTalairach) and visualized using Caret software (53) (<u>http://www.nitrc.org/projects/caret/</u>, v5.65). Overlap of the gastric network with cytoarchitectonic subdivisions of primary and secondary somatosensory cortices was determined with the anatomy toolbox for SPM (54–56).

### Data availability statement

Unthresholded t maps of empirical vs chance PLV comparisons (intermediate step for Fig. 2a), mask of significant clusters (Fig. 2a) and average phase-locking angle of each significant cluster (Fig. 4a) are available at Neurovault (57) at the following address: <u>http://neurovault.org/collections/GMHHGEXA/</u>

### Code availability statement

The custom code used for this article can be accessed online at the following address: <u>https://github.com/irebollo/stomach_brain_Scripts</u>.

## Acknowledgments

This work was supported by funding from the European Research Council (ERC) under the European Union’s Horizon 2020 research and innovation program (grant agreement No 670325) to CTB, as well as from ANR-10-LABX-0087 IEC and ANR-10-IDEX-0001-02 PSL*. I.R. is supported by a grant from DIM Cerveau et Pensée Région Ile de France and by the Fondation Bettencourt-Schueller

## Authors contributions

I.R. and C.T.-B. designed the experiment; I.R., A.-D.D, and B.B. acquired the data; I.R. and A.-D.D. preprocessed the data; and I.R. and C.T.-B. analyzed the data and wrote the paper.

